# Genotypic variation in root distribution changes and physiological responses of sugarcane induced by drought stress

**DOI:** 10.1101/503912

**Authors:** Jariya Namwongsa, Nuntawoot Jongrungklang, Patcharin Songsri

## Abstract

Drought is an important factor reducing yield and quality of sugarcane. Root growth and physiological traits are important for maximizing water uptake to improve drought resistance. This study compared the root, shoot and physiological traits under drought stress (DS) and well-watered (WW) conditions of various sugarcane varieties grown in rhizoboxes in a greenhouse at the Field Crops Research Station of Khon Kaen University, Khon Kaen, Thailand. Data were recorded for the root, shoot and physiological traits (relative water content, stomatal conductance, SPAD chlorophyll meter reading and chlorophyll fluorescence) at 90 days after transplanting. Root samples were recovered from 11 soil layers at 10 cm intervals from the top to the bottom of the rhizobox, for root length and root dry weight measurements. Drought was imposed on sugarcane at early growth stages altered the root distribution patterns, and differences were evident among the sugarcane genotypes. The sugarcane genotypes adapted to water stress by increasing root length into deeper soil layers. Drought led to increased total root length in KK3, MPT06-166, K88-92, CP38-22, Kps01-12 and KPK98-40. Root lengths and stomatal conductance were positively correlated under WW and DS conditions. Root distribution in the lower soil layers and the percentage of root distribution were higher than those under well-watered conditions. The knowledge gained from this study will aid parental selection in sugarcane breeding programs for drought resistance as the findings strongly suggest that the physiological modification in the root system is a useful drought-resistant mechanism.

## Introduction

Sugarcane is the world’s largest crop by production quantity, and it is cultivated in about 120 countries in the tropical region with the global harvest now exceeding 175 million tons a year (FAO, 2016). Sugarcane is used primarily for sugar production and an efficient crop for other products, such as electricity, bioethanol and fertilizer (Unica, 2008). Despite increasing consumer demands for sugar, the cane yield and sugar yield in production systems are still low due to diseases, insect infestations and drought. Early season drought and mid-season drought can reduce plant growth, resulting in plant stunting and restriction of tillering, leading to vacant and low millable stalk and losses in both cane yield and sugar yield (Dinh et al., 2017). Drought stress (DS) can cause yield losses of up to 60% (Robertson et al., 1999; Gentile et al., 2015). A drought-resistant sugarcane cultivar could maintain yield under rainfed and DS conditions. However, an understanding on the drought-resistant mechanisms is a major challenge in sugarcane breeding programs as drought resistance is inherited genetically and closely related to physiological characteristics.

Common physiological traits including leaf area, stomatal conductance, chlorophyll content, relative water content (RWC), photosystem II (PSII) photosynthesis efficiency and photosynthetic rate have been used to improve drought resistance in sugarcane breeding (Silva et al., 2008). Moreover, reduction in stomatal conductance (to reduce water loss) and an increase in density and deep root traits (to increase water uptake ability) have been reported as the mechanisms for adaptation in sugarcane to maintain water status in the plant under water stress conditions (Wasson et al., 2012). Therefore, DS causes a decrease in stomatal conductivity to reduce water loss in the leaf. The resulting carbon dioxide entering through the stomata is also reduced as a result of a reduction in photosynthesis, which may also lead to low sugarcane yields. The mechanism of drought avoidance associated with root characters is to search for water in the soil layers. Under well-watered conditions, most root systems of sugarcane are in the upper soil layers (Smith et al., 2005), and a decrease in the moisture content of the soil surface activates growth of the roots in the lower soil layers. The adapted root system acts to promote water absorption, thereby maintaining the water balance in the plant, and the adaptation of root system increases the amount of transpiration, which helps the sugarcane to avoid the danger of dehydration.

Studies of root traits such as physiological parameters can be useful to select sugarcane genotypes for drought avoidance mechanism. The standard method used in previous investigations is to evaluate root growth and distribution pattern of sugarcane under the natural conditions provided by field experiments. However, among the methods for collecting root data, such as drawings, monoliths and auger methods, considerable time, labor and cost are involved because the actual growth of the sugarcane roots is not visible. Although greenhouse experiment is an indirect and cost-effective means of studying root distribution, evaluation the plant root growth is limited to the early growth stage. Studies of root systems in the rhizobox are mostly done with young plants or annual plant species. The use of split-root systems to monitor the effect of root distributions on the development of the root system is very interesting (Neumann et al., 2009). The rhizobox is effective for displaying the characteristics of the root distribution and this method has been used in many plant species such as peanuts (Thangthong et al., 2016; Thangthong et al., 2017) and Jerusalem artichoke (Puangbut et al., 2018) in which different varieties showed differences in the root distribution patterns. The information on the changes in root distribution patterns under well-water condition and drought condition for characterization of sugarcane genotypes is still lacking.

The aim of this study was to investigate root distribution and physiological responses under WW and DS conditions for various sugarcane varieties grown in rhizoboxes. The data were evaluated to understand the response of root distribution and physiological characteristics under WW and DS conditions. The information obtained in this study is necessary for further experiments and may be applied for selection of sugarcane varieties for drought resistance.

## Materials and methods

### Plant materials and rhizobox preparation

The experiment was conducted in rhizoboxes in a greenhouse at the Field Crops Research Station of Khon Kaen University, Khon Kaen, Thailand (16°28’N, 102°48’E, 200 m above sea level) during 21 June to 17 September 2016. The 13 sugarcane genotypes consisting of Yasawa, MPT03-320, PR3067, KK3, MPT06-166, K88-92, CP38-22, UT5, Kps01-12, KPK98-40, F152, BO14 and Nco382 were selected by screening for differences in total root length, using a small pot experiment (data not presented).

In this study, root distribution and root architecture were investigated using the modified box–pinboard method (needle-board) (Fig. 1). The detailed method was clearly described in the previous studies (Thangthong et al., 2016; Thangthong et al., 2017), and the method was briefly described herein.

**Fig. 1.**
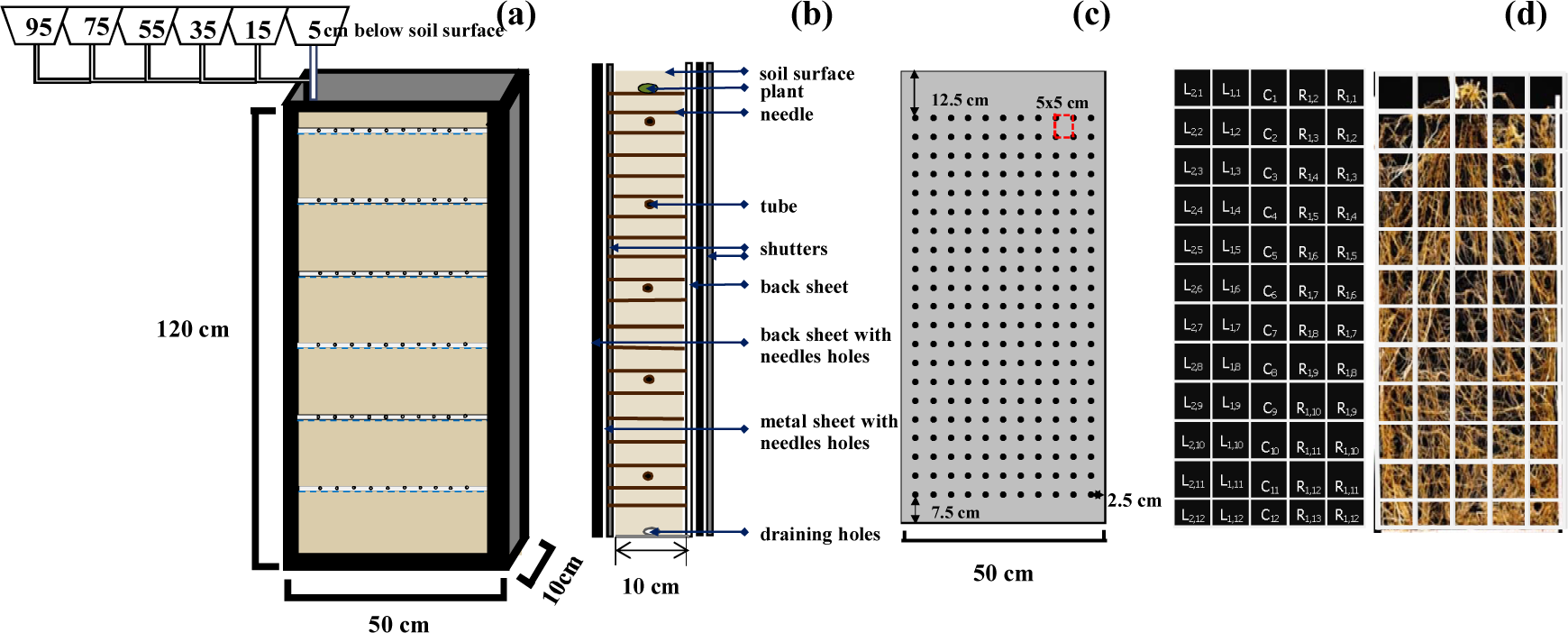
Diagrammatic representation and dimension of rhizobox with the positions of six tubes of irrigation (a), the section showing different elements of the system (b), spacing of needles at backside of rhizoboxes (c), and size of the square units of 10 cm × 10 cm in which the roots were observed and the image of sugarcane root system that was grown in rhizobox at 90 days after planting (d)

The dimension of the rhizoboxes was 10×50×120 cm, the boxes were filled uniformly with dry soil to reach the height of 115 cm. The soil was divided into 11 layers from the bottom of the boxes to the top of the boxes. The boxes had a needle grid with a spacing of 5×5 cm at the back of the boxes to held root at the original positions after washing. The boxes also had a transparent window at the front of the boxes for visually observing root growth and taking photographs of root growth.

A sugarcane sett with a germinated single bud was plated at the center of the boxes at 5 cm below the soil surface. The boxes were wrapped with a black sheet at all sides of the boxes and then the boxes were wrapped again with an aluminum foil. The front side of the boxes could be opened to see the transparent window. This method has been used to study root systems in some plant species, such as peanut (Thangthong et al., 2016; Thangthong et al., 2017) and Jerusalem artichoke (Puangbut et al., 2018).

### Irrigation treatments

Irrigated water was supplied to the rhizoboxes through tube irrigation system. Six tubes for each rhizoboxes were installed at 5, 15, 35, 55, 75 and 95 cm below the soil surface (Fig. 1a) and, before transplanting, the water was supplied at field capacity (FC) to all experimental units (rhizobox) throughout the boxes. At 10 days after transplanting (DAT), water was provided at the soil surface of the two treatments based on the water requirement of sugarcane for the uniformity of sugarcane sett germination.

Two water regimes consisting of well-watered (WW) level and drought stress (DS) level were created. At 10 DAT, WW treatment was supplied to the boxes through three upper tubes at 5, 15 and 35 cm below the soil surface at FC level from initiation of the experiment until 45 DAT. At 45 DAT, WW treatment was supplied to the boxes at FC level until the end of the experiment through six tubes mounted at 5, 15, 35, 55, 75 and 95 cm below the soil surface.

DS treatment was supplied to the boxes at FC level from initiation of the experiment until 30 DAT through three tubes mounted at 5, 15, 35 cm below the soil surface, and then the irrigated water was reduced to half of FC level until 45 DAT. From 45 DAT to the end of the experiment, DS was supplied to the boxes at half of the FC level through three tubes mounted at 55, 75 and 95 cm at the lower soil layers. The soil moisture reduction was simulated similar to that under field condition to create higher soil moisture in the lower soil layers than in the upper soil layers.

The water requirement of sugarcane was calculated daily, as the sum of water loss through transpiration and soil evaporation based on the crop water requirement (*ET*_crop_) (Doorenbos and Pruitt, 1992; Jangpromma et al., 2012):

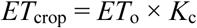

where *ET*_crop_ = crop water requirement (mm day^-1^); *ET*_o_ = evapotranspiration of a reference plant under specified conditions, calculated by the class A pan evaporation method (mm day^-1^), and *K*_c_ = the crop water requirement coefficient for sugarcane.

### Plant management

Before transplanting, each sugarcane genotype in the rhizoboxes was subjected to a curing process for 5 days until the buds and root primordial (0.5 cm) were germinated. Fertilizer grade 15-15-15 was applied at 1.56 g per rhizobox at 1 DAT. Fertilizer grade 46-0-0 was applied at 1.56 g per rhizobox at 60 DAT. Soil moisture contents at FC (13%) and permanent wilting point (4.3%) were determined by the pressure plate method.

### Data collection

#### Soil moisture content

Soil moisture content was measured gravimetrically, using a micro-auger at 14, 28, 45, 60 and 90 DAT. Soil moisture content was collected at soil depths of 10 cm (14 DAT); 10 and 25 cm (28 and 45 DAT); 10, 25, 45, 65 and 85 cm (60 DAT); and 10, 25, 45, 65, 85 and 105 cm (90 DAT), respectively. Soil moisture content for each rhizobox was calculated as follows:

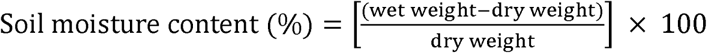

### Root characteristics

Root traits were measured at 90 DAT. The rhizoboxes were carefully washed with tap water to clean the root samples. The root arrangement in the soil was fixed by needles and the needles on the back of the boxes were removed. Thereafter, two procedures were used to determine root traits: (i) taking photographs using a CanonEOS5D Mark IV24-70 f2.8 (Canon Ltd., Tokyo, Japan), and (ii) root scanning by using an Epson (Perfection V700, Japan) for analysis of the root length. The photographs showed the root distribution patterns of the whole root system on a black sheet with a scale bar. Roots were separated into square sections taken from the left, center and right columns. Root sample of each rhizobox was divided into 11 soil layers at 10 cm intervals from the top to the bottom of the box (Fig. 1d).

For root scanning, the sample was separated into square unit sizes of 10 cm × 10 cm (Fig. 1d). Root length was analyzed by the WinRHIZO program (WinRHIZO Pro(s) V.2004a, Regent Instruments Inc. Canada) for the root distribution patterns. Root lengths in the upper soil layers of 0–10, 10–20, 20–30 cm were combined into a single 0–30 cm layer, whereas root traits at the lower soil layers were combined to form a single layer of 30–110 cm.

### Root and shoot dry weight

At 90 DAT, plant shoots were cut at the soil surface and shoot fresh weight was recorded. The samples were then oven-dried at 80 °C for 48 h and shoot dry weight was recorded. After scanning for root length measurement, the root samples were oven-dried at 80 °C for 48 h and root dry weight was recorded.

### Physiological characteristics

SPAD chlorophyll meter reading (SCMR), chlorophyll fluorescence, stomatal conductance and RWC were recorded in each rhizobox at 90 DAT. All characteristics were recorded during 09.00–12.00 h. SCMR was recorded on the second or third fully-expanded leaf from the top of the main stalk using a SPAD-502 chlorophyll meter (Minolta SPAD-502 meter, Tokyo, Japan).

The same leaf samples were used for recording chlorophyll fluorescence using a chlorophyll fluorescence meter (MINI-PAM-2000, Heinz Walz GmbH, Germany). The leaf samples were stored under dark for 30 min, and chlorophyll fluorescence was recorded using leaf clips (FL-DC, Opti-Science, Wetzlar, Germany). Chlorophyll fluorescence was determined according to the method Maxwell and Johnson 2000 described previously to quantify the level of drought-induced photo-inhibition.

Stomatal conductance was measured on the intact leaves. The second or third fully-expanded leaf from the top of the main stalk was used for measurement of the trait using a porometer (model AP4, Delta-T Devices, Cambridge, UK).

The same leaf samples for measurement of stomatal conductance were used for measurement of relative water content (RWC). The samples were harvested from the plants and leaf fresh weight was recorded. The leaf samples were placed in deionized water for 24 h at room temperature and leaf turgid weight was recorded. Leaf dry weight was measured after oven-drying at 80 °C for 48 h (Silva et al., 2007). RWC was calculated using the formula below:

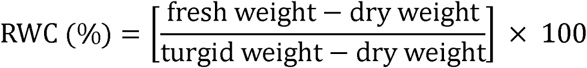

## Results

### Soil moisture content

Sugarcane varieties grown under well-watered condition and drought stress conditions were different for soil moisture content from 45 to 90 DAT (Fig. 2a). Water regimes were different in soil moisture content in the soil depth of 10–85 cm at 60 DAT (Fig. 2b) and 10– 105 cm at 90 DAT (Fig. 2c).

**Fig. 2.**
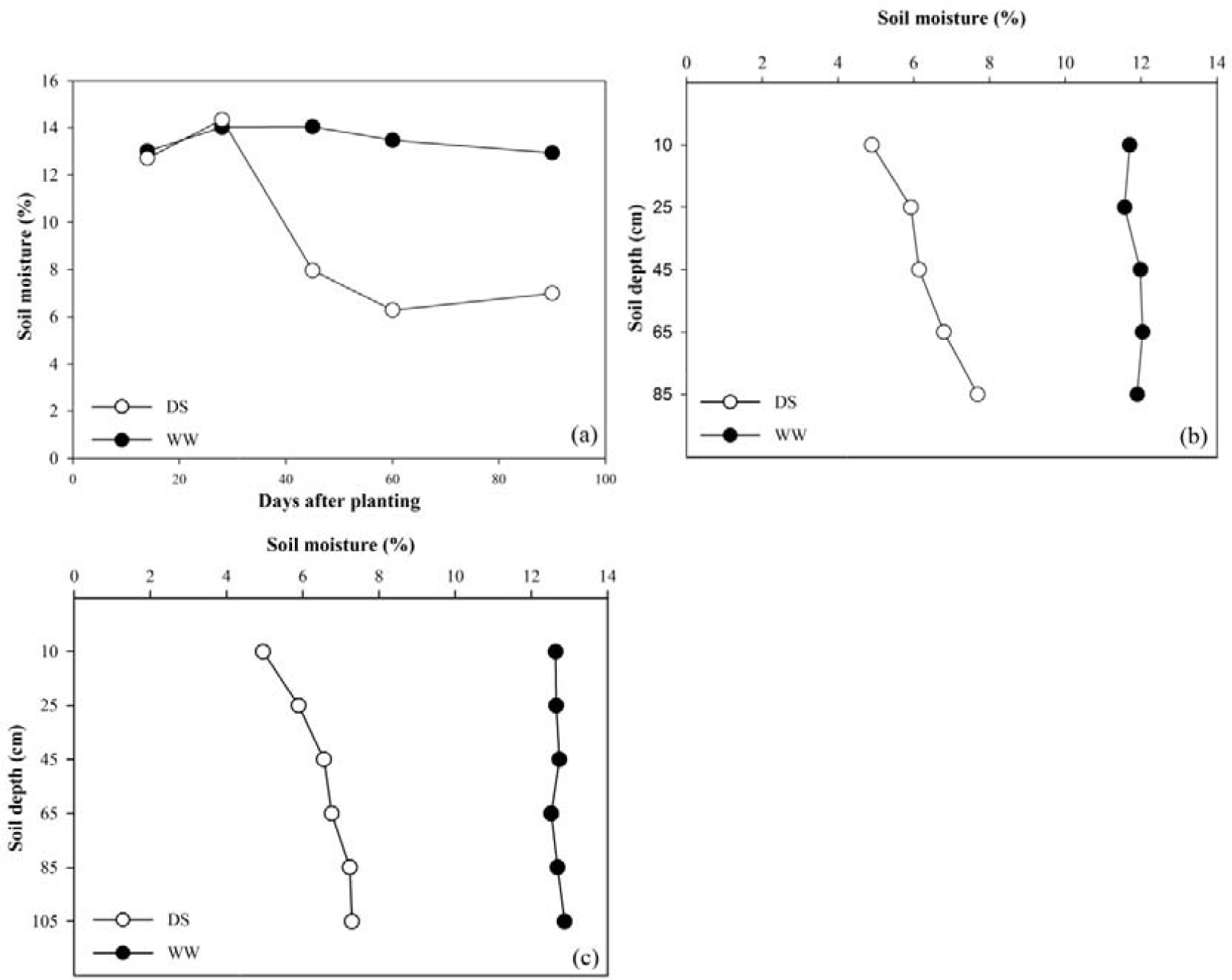
Soil moisture content (%) of drought stress (DS) and well-watered (WW) treatments at 14, 28, 45, 60 and 90 days after planting (a) and in different soil layers 10, 25, 45, 65 and 85 cm at 60 DAP (b) and 10, 25, 45, 65, 85 and 105 cm at 90 DAP (c)

### Root distribution pattern of sugarcane

Images of the root distribution patterns of all 13 sugarcane genotypes grown in the rhizoboxes (Fig. 3) and captured at 90 DAT revealed the root distribution patterns of all the genotypes under WW condition and DS condition. Also, superficial roots appeared mostly in the upper soil layer (Fig. 3, 4).

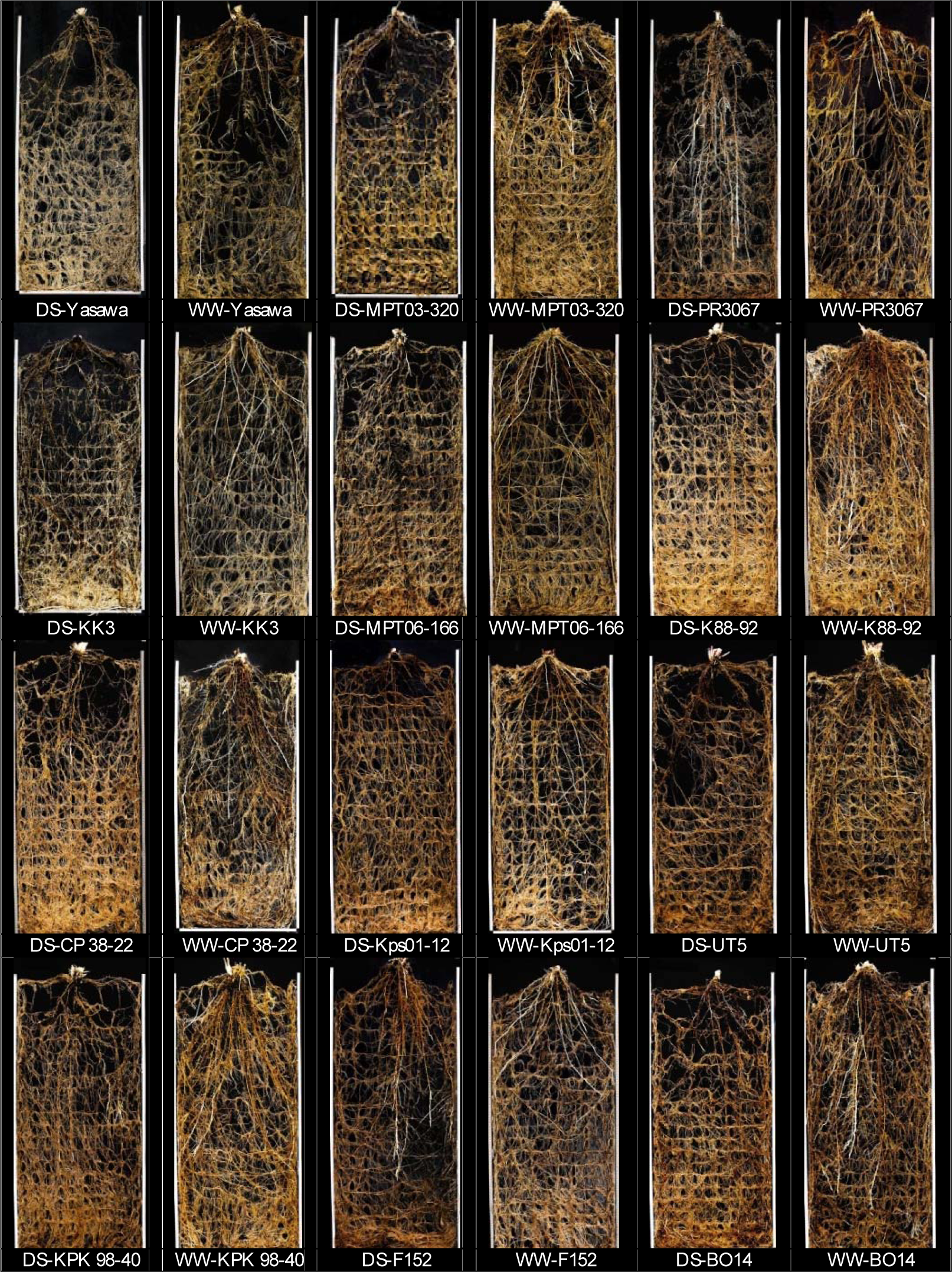
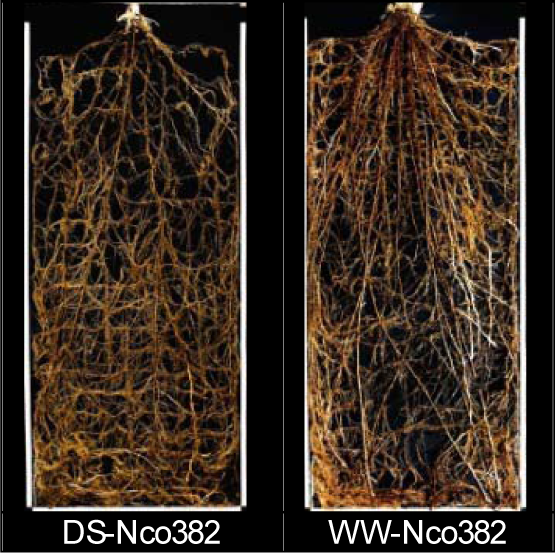

**Fig. 4.**
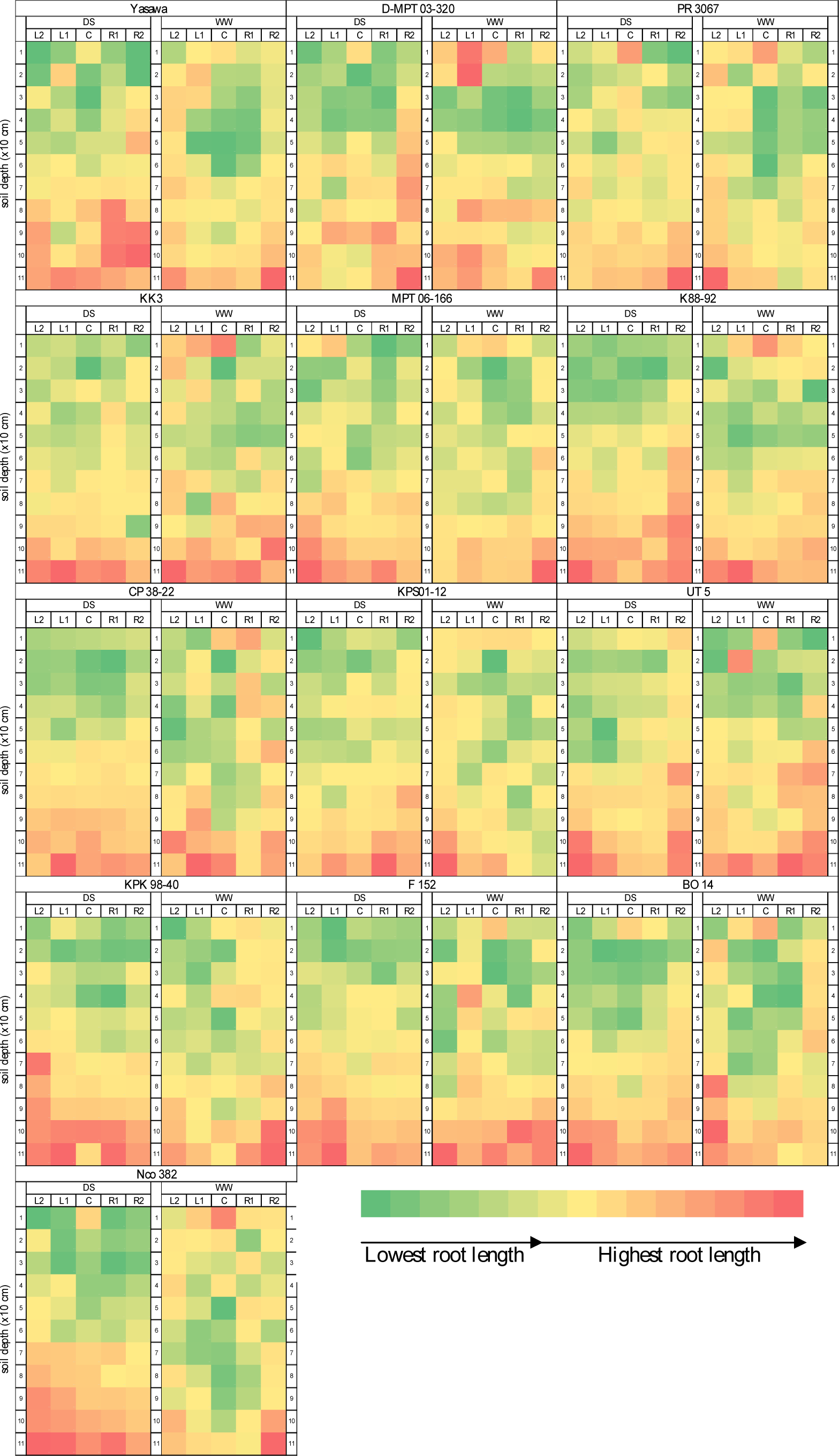
Graphical presentation describing root length distribution patterns of 13 sugarcane genotypes grown in rhizobox under drought stress (DS) and well-watered WW conditions at 90 days after planting

Roots of sugarcane genotypes were reduced in upper soil layers (0–30 cm) in response to drought stress, but the roots in the lower soil layers (below 30 cm) were increased. KK3, K88-92, CP38-22, KPK98-40, BO14 and Nco382 increased new roots (white color) the lower soil layers, whereas PR3067 and F152 increased buttress roots the lower soil layers to take up water and minerals from deep soil layers especially under drought stress (Fig. 3).

KK3, K88-92, CP38-22, Kps01-12 and BO14 increases buttress root the lower soil layers (Fig. 3, 4). Under well-watered condition, K88-92, KPK98-40, BO14 and Nco382WW developed superficial roots in soil surface (Fig. 3). Under well-watered condition, CP38-22 and KPK98-40 could maintain high root growth as indicated by high root length. Under drought stress condition, they reduced root growth in the upper soil layers and increased root growth in the lower soil layers.

Nco382 had higher root length under well-watered condition than under drought condition as root length was reduced under drought stress. (Fig. 3). KK3 had higher root length in lower soil layers under drought stress condition than under well-watered condition. Kps01-12 under drought stress could maintain high root length both in upper soil layers and lower soil layers. K88-92 and BO14 under well-watered condition had high root length both in upper soil layers and lower soil layers. Under drought stress condition, these genotypes reduced root length in upper soil layers and increased root length in lower soil layers. Under drought stress condition, MPT03-320 and MPT06-166 reduced root length in upper soil layers to maintain root growth in lower soil layers.

DS and WW treatments were compared for root length and root distribution at 90 DAT (Fig. 5). For each of the 11 soil layers (at 10 cm intervals from the top to the bottom of the rhizobox), the root length increased at initiation stage of water with-holding up to 90 DAT. The sugarcane crops grown under drought treatment and well-watered treatment were significantly different for root length at 90 DAT. The percentage of root length in the upper soil (0–30 cm) was lower than that in the lower soil (below 30 cm) (Fig. 5). Root lengths of most sugarcane genotypes grown under DS condition were longer than those under WW condition in most of the soil layers (Fig. 5) except for MPT03-320, PR3067 and UT5 genotypes, which had the opposite trend (Fig. 5). All sugarcane genotypes grown under drought conditions had higher percentage of root length than did these genotypes grown under well-watered conditions. Most sugarcane genotypes grown under drought conditions had lower root percentage in the upper soil layers than did these genotypes grown under well-watered conditions except for UT5, which had similar percentage of root length both under drought conditions and well-watered conditions (Fig. 5). However, the patterns of root growth between DS and WW treatments were rather different in all 13 genotypes, and the sugarcane genotypes grown under drought conditions had higher percentages of roots and root length in the lower soil layers than did these genotypes grown under well-watered conditions. The results indicated that sugarcane genotypes grown under drought conditions distributed a greater proportion of roots in the lower soil layers than did these genotypes grown under well-irrigated.

**Fig. 5.**
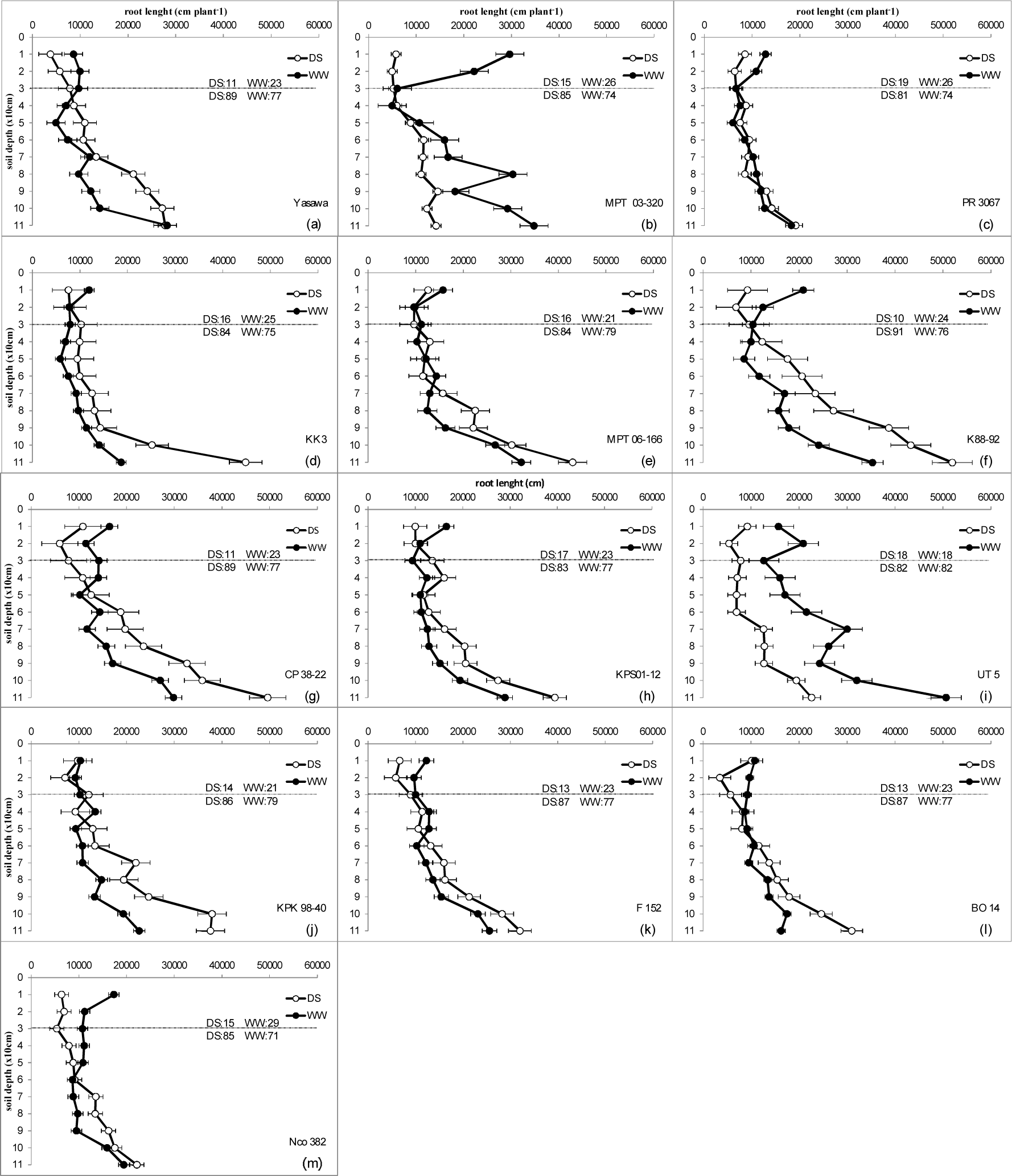
Root length distribution of sugarcane under drought stress (DS) and well-watered (WW) of 11 (10 cm interval) layers divided into two soil layers defined previously as upper (0–30 cm of soil depth) and lower (30–120 cm of soil depth) evaluated at 90 days after planting.

### Responses of root characteristics to water regimes

Sugarcane genotypes were significantly different for total root length. Drought stress increased total root length in KK3, MPT06-166, K88-92, CP38-22, Kps01-12 and KPK98-40. It reduced root length in MPT03-320 and UT5, whereas Yasawa, PR3067, F152, BO14 and Nco382 were not significantly affected by drought stress (Fig. 6a).

**Fig. 6.**
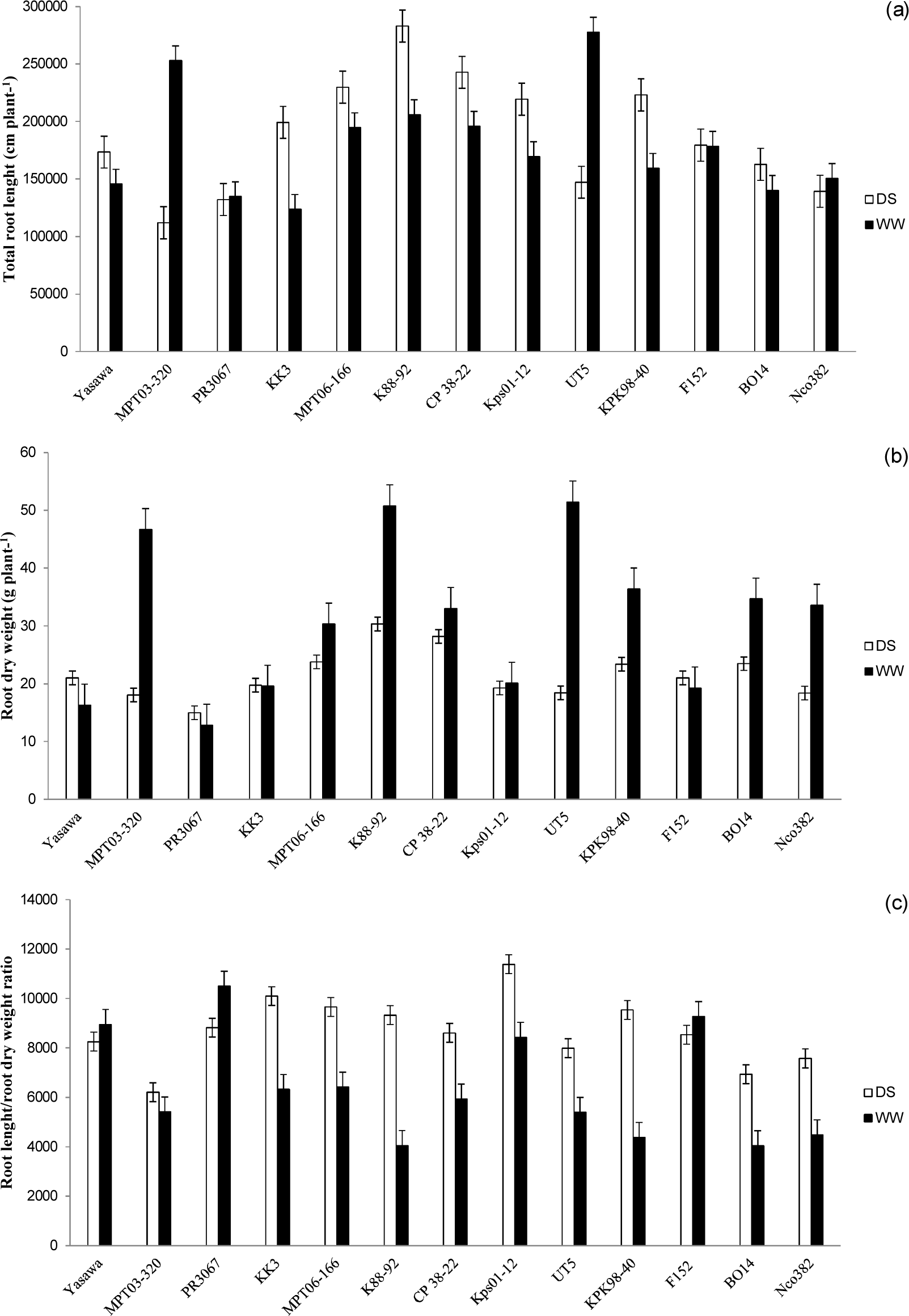
Total root length (a), root dry weight (b) and root length/root dry weight ratio (c) of 13 sugarcane genotypes grown in rhizobox under drought stress (DS) and well-watered (WW) conditions

Under drought conditions, KK3 and Kps01-12 could maintain root dry weight (root size) similar to that under well-watered conditions but they did increase root length. MPT06-166, K88-92, CP38-22, KPK98-40 and BO14 decreased root dry weight but increased root length. MPT03-320, UT5 and Nco382 displayed a decrease in both root dry weight and root length. For the remaining genotypes, the results were not significantly different between the two water regimes (Fig. 6b).

MPT03-320, KK3, MPT06-166, K88-92, CP38-22, Kps01-12, UT5, KPK98-40, BO14 and Nco382 had an increased root length-to-root dry weight ratio under DS conditions. The opposite trend was observed for Yasawa, PR3067 and F152 (Fig. 6c).

### Biomass, shoot dry weight and root-to-shoot ratio

Sugarcane genotypes were significantly different for biomass, shoot dry weight and root-to-shoot ratio under drought stress conditions and well-watered conditions (Fig. 7a–c). Under well-watered conditions (Fig. 7a), MPT03-320, K88-92, CP38-22, UT5, BO14 and Nco382 had high biomass, KPK98-40 had moderate biomass, and Yasawa, PR3067, KK3, MPT06-166, Kps01-12 and F152 had low biomass. Under drought stress conditions, MPT03-320, K88-92, CP38-22, UT5 and Nco382 reduced biomass, but PR3067 increased biomass, whereas Yasawa, KK3, MPT06-166, Kps01-12, KPK98-40, F152 and BO14 had similar biomass both under drought stress conditions and well-watered conditions.

**Fig. 7.**
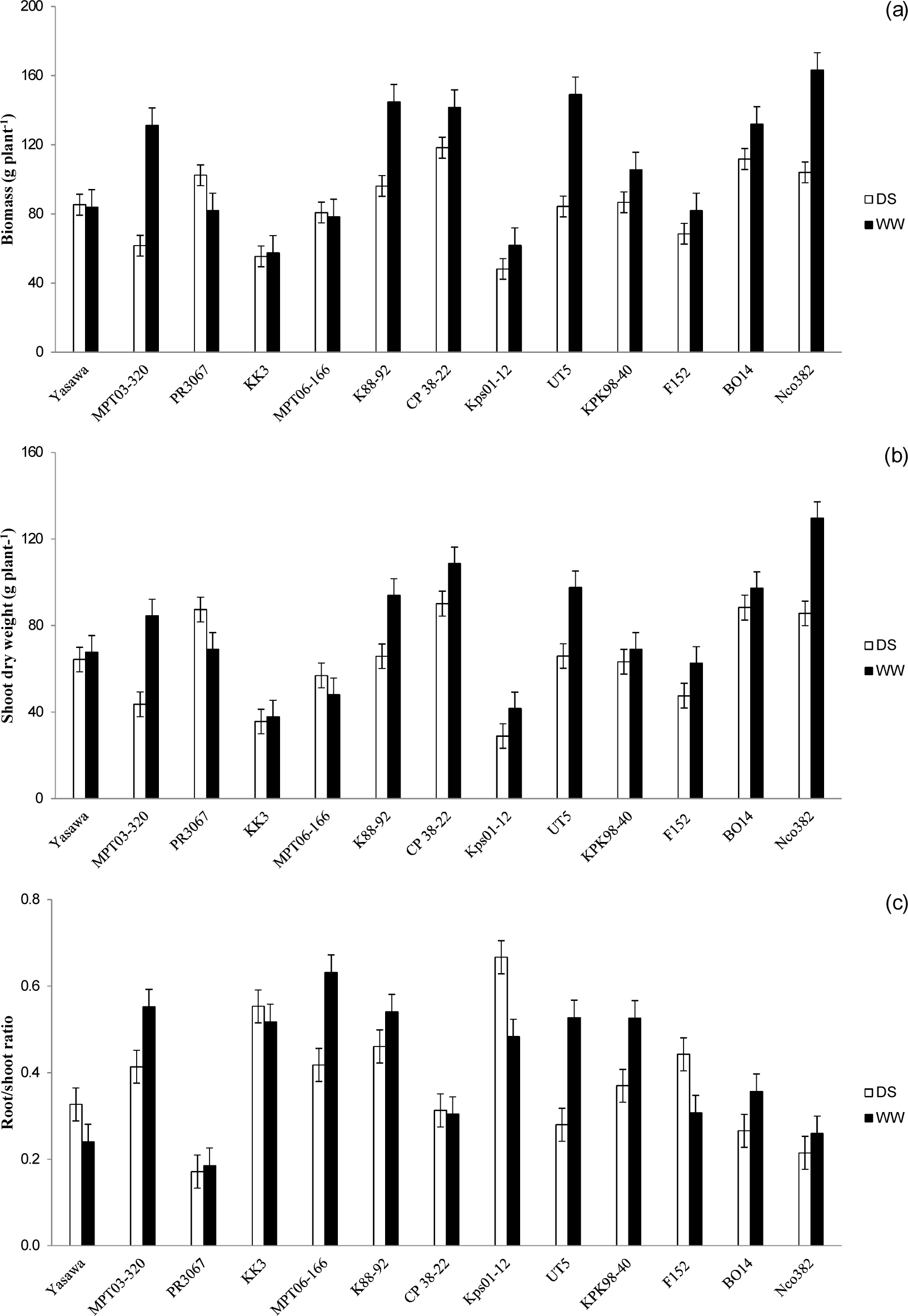
Biomass (a), shoot dry weight (b) and root/shoot ratio (c) of 13 sugarcane genotypes grown in rhizobox under drought stress (DS) and well-watered (WW) conditions

MPT03-320, K88-92, CP38-22, Kps01-12, UT5 and Nco382 reduced shoot dry weight and PR3067 increased shoot weight, whereas Yasawa, KK3, MPT06-166, KPK98-40, F152 and BO14 had similar shoot dry weight both under drought stress conditions and well-watered conditions (Fig. 7b). Similar to biomass and shoot dry weight, under drought stress conditions, MPT03-320, MPT06-166, K88-92, UT5, KPK98-40 and BO14) reduced root-to-shoot ratio, Yasawa, Kps01-12 and F152 increased root-to-shoot ratio, whereas PR3067, KK3, CP38-22 and Nco382 had similar root-to-shoot ratio both under drought stress conditions and well-watered conditions (Fig. 7c).

### Relationship of root traits and above-ground traits between stress and non-stress

The correlations between sugarcane varieties grown under drought stress conditions and well-watered conditions were positive and significant (P≤0.05 and 0.01) for shoot dry weight, biomass and the root-to-shoot ratio (*r* = 0.74, 0.63 and 0.57, respectively) (Fig. 8c, 8f, 8d). However, the correlations between sugarcane varieties grown under drought conditions and well-watered conditions were not significant for total root length, root dry weight and root length-to-root dry weight ratio (Fig. 8a, 8b, 8e).

**Fig. 8.**
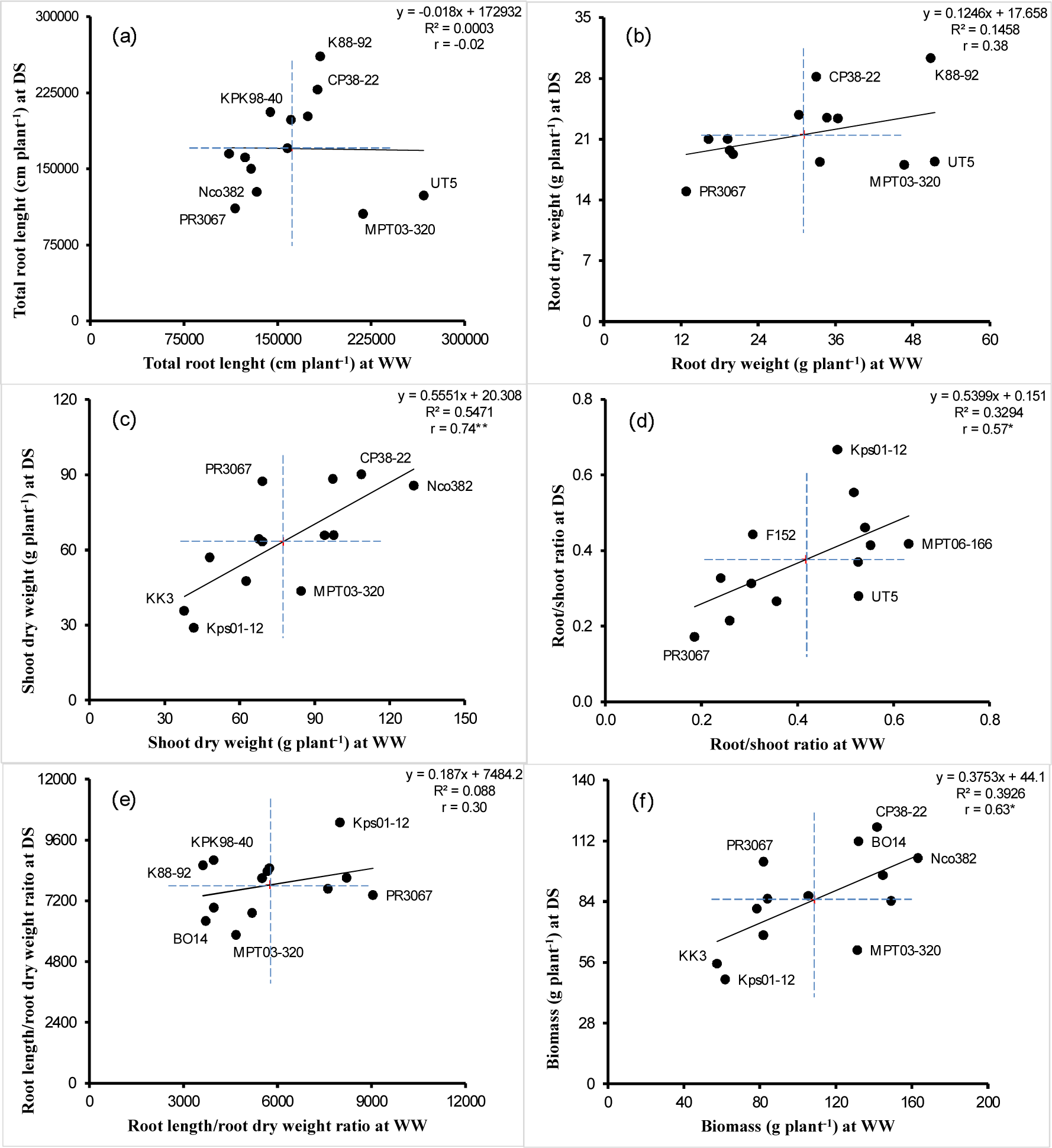
Relationships between 13 sugarcane genotypes grown in rhizobox between under drought stress (DS) and well-watered (WW) conditions for root length (a), root dry weight (b), shoot dry weight (c), root/shoot ratio (d), root length/root dry weight ratio (e) and biomass (f)

Sugarcane genotypes were significantly different for root length under drought stress conditions. K88-92, CP38-22 and MPT06-166 had high root length under well-watered conditions (high potential) or both under well-watered conditions and drought stress conditions. KPK98-40, Kps01-12, KK3 and F152 increased root length under drought stress conditions, MPT03-320 and UT5 reduced root length under drought stress conditions, whereas Yasawa, BO14, Nco382 and PR3067 had similar root length both under drought stress conditions and well-watered conditions.

### Relationships between root and physiological traits under stress and non-stress conditions

The correlations between stomatal conductance and total root length were positive and significant both under well-watered conditions (*r* = 0.81, P≤ 0.01) and drought stress conditions (*r* = 0.75, P≤ 0.01) (Fig. 9c, 10c), whereas the correlations between other physiological traits and root traits were not significant. Under drought stress conditions and well-watered conditions, K88-92, CP38-22 and MPT06-166 with high root lengths were strongly associated with high stomatal conductance (Fig. 9c, 10c).

Relative water content indicates the plant water status. Under drought stress conditions, K88-92, CP38-22 and MPT06-166 had low relative water content (Fig. 9b). The sugarcane genotypes grown under drought stress conditions and under well water conditions at 90 DAT were significantly different for relative water content (Fig. 9b, 10b). The differences in relative water content between drought stressed crop and well watered crop would be due to difference in soil moisture content. Relative water content was appropriate for evaluation of plant water status because the traits was associated soil moisture content.

**Fig. 9.**
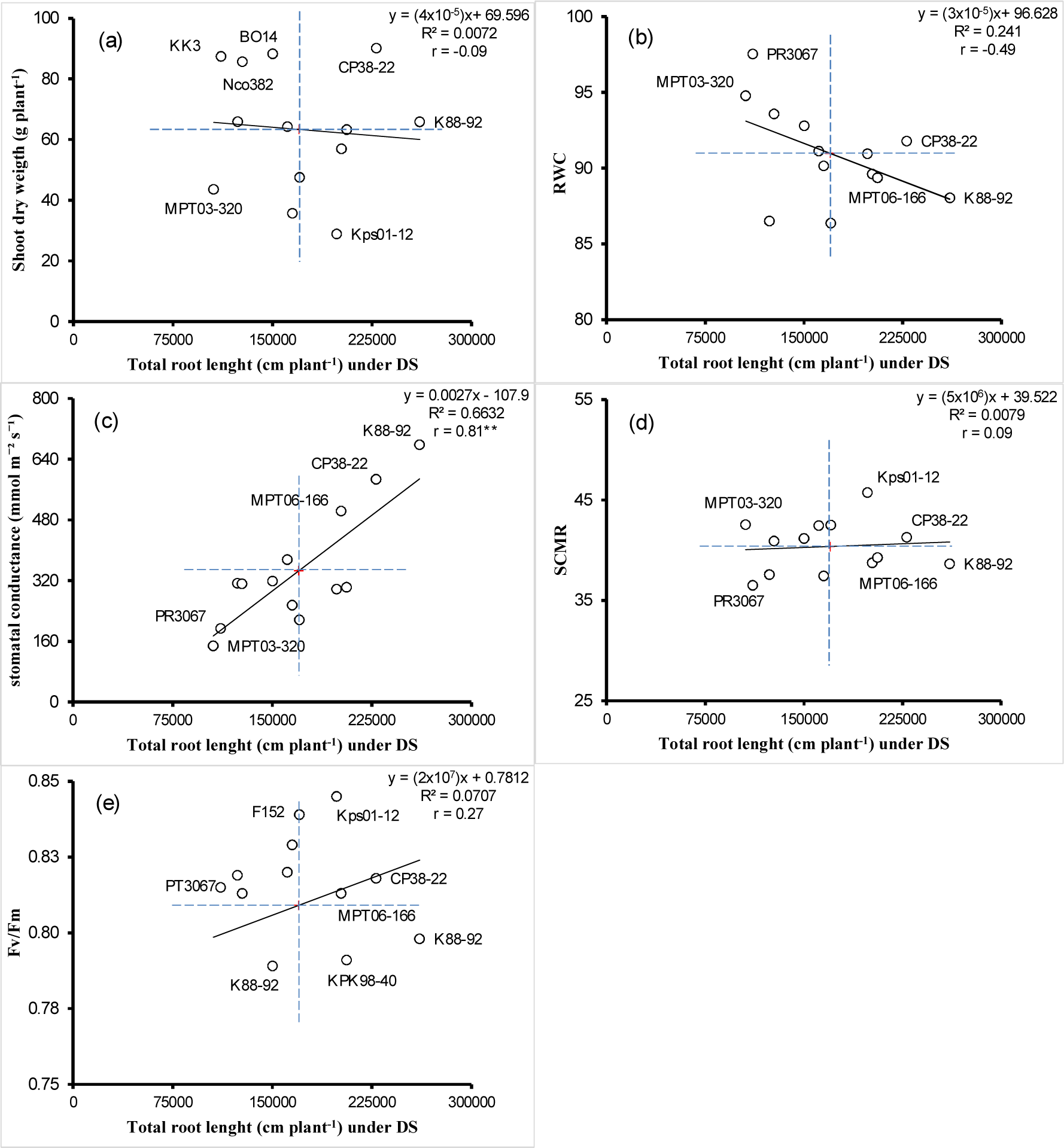
Relationships between total root length and biomass (a), relative water content (RWC) (b), stomatal conductance (c), SPAD chlorophyll meter reading (SCMR) (d) and chlorophyll fluorescence (e) of 13 sugarcane genotypes grown under drought stress (DS) conditions

For the genotypes (K88-92, MPT03-320, CP38-22, MPT06-166 and UT5) with high total root length per plant under well-watered conditions, total root length was associated with stomatal conductance (Fig. 10c), whereas total root length was not associated with relative water content (Fig. 10b). Sugarcane genotypes were significantly different for total root length under drought stress conditions. The correlation coefficients between root length and shoot dry weight (Fig. 9a, 10a), RWC (Fig. 9b, 10b), SCMR (Fig. 9d, 10d) and chlorophyll fluorescence (Fig. 9e, 10e) were not significant.

**Fig. 10.**
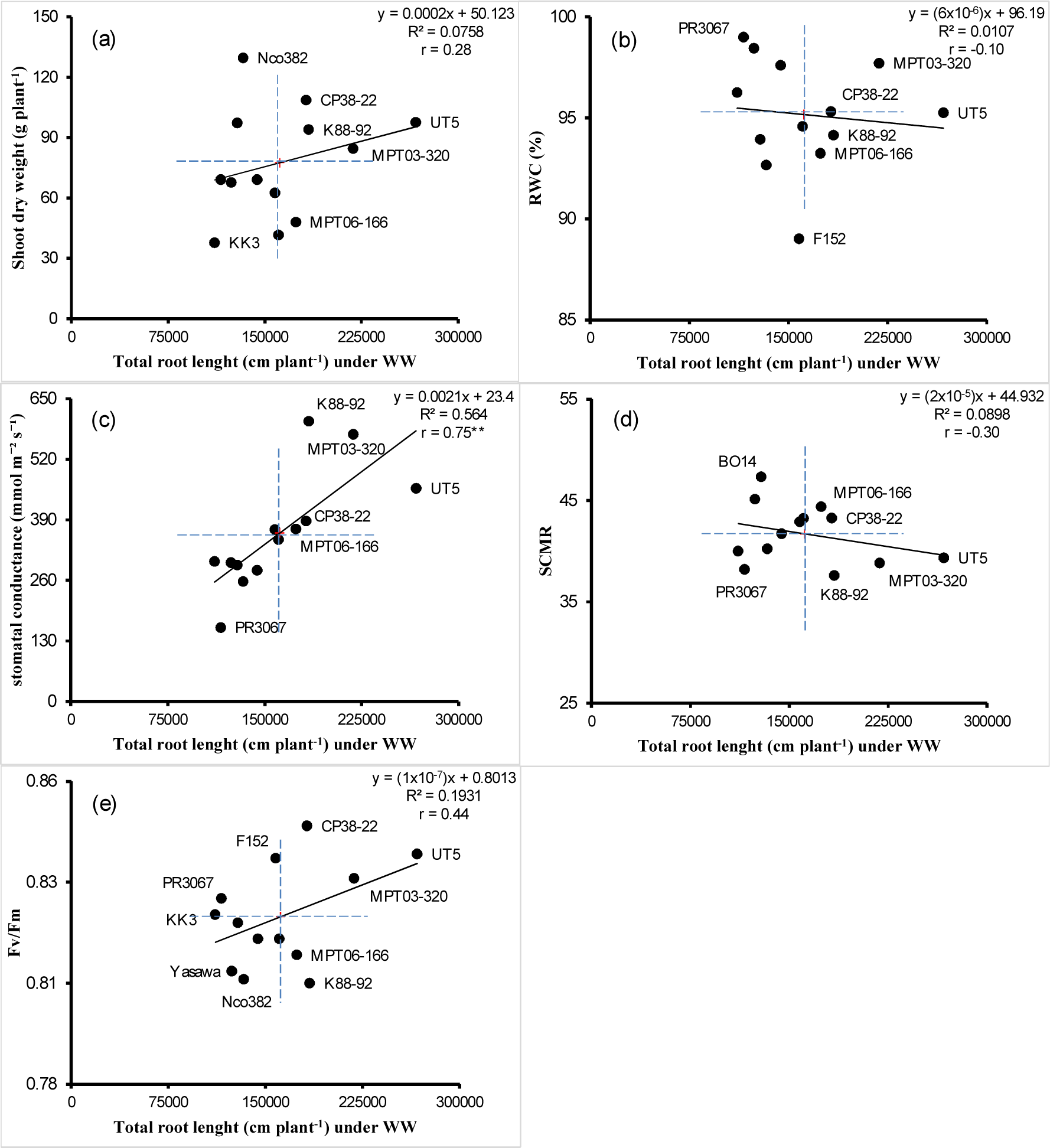
Relationships between total root length and biomass (a), relative water content (RWC) (b), stomatal conductance (c), SPAD chlorophyll meter reading (SCMR) (d) and chlorophyll fluorescence (e) of 13 sugarcane genotypes grown under well-watered (WW)

## Discussion

### Root development for sugarcane growth

Root growth and distribution under diverse environments are important for predicting the responses of the plant to changes in soils and environments. Drought tolerance strategies regularly include the development of expanded or deep root structures and other physiological functions such as oxidative stress protection and a decrease in transpiration and osmoregulation (Pirnajmedin et al., 2015). The main findings of this study are that the water-limited condition changed the root distribution patterns of the different sugarcane genotypes investigated. The plant roots grew into the deeper soil layers in response to the decrease in soil water. The differential responses of sugarcane genotypes to drought stress for root traits and other physiological traits may be depend on genetic control. Under water-limited conditions, the morphology of root structures is essential in accessing nutrients and soil water (Smith et al., 2005). Rapid development and suitable distribution of the sugarcane root system at deeper soil layers are essential to curtailing the adverse consequence of these dry periods on yield. Under well-watered conditions, root density was high and uniform throughout the soil layers (Kato and Okami, 2011). Under water-limit condition, plant invests in root growth higher than in shoot growth to take up more water. Under water-deficit condition, translocation of assimilates to roots was higher than to shoots (Azhiri-Sigari et al., 2000). In the course of the early growth stages of the water-limited condition, progressive accumulation of root dry matter was on the expense of shoot growth, and the plants with adaptation to dry condition had higher root/shoot ratio. Once a decrease in the soil moisture content is detected, the roots must expand their distribution patterns and elongate into deeper soil layers for extracting and engaging a larger soil volume for water. As soil moisture at the soil surface and in the top soil profile was diminished under water deficit stress, the roots removed more water at the deeper profile. A deep root scheme is helpful for extracting water form substantial soil depths (Kavar et al., 2007). This root system characteristic is an important consequence to soil drying and allows some roots to continue elongation under a water deficit to search more water. The distribution of the root schemes depended strongly on the soil moisture of the deeper soil layer.

### Root characteristic under water deficit

The water deficit might adversely affect root growth in the upper soil layers as the upper soil layers are dryer than the lower soil layers. Root growth in upper soil layers is then limited by the drying soil, whereas root growth in the lower soil layers is still continued in response to soil moisture. Root response to soil moisture in the lower soil is an important character to enhance water extraction from the deeper soil and improve the plant’s potentiality to resume growth during drought. It has been proposed that breeding for a narrow xylem vessel in the seminal roots of wheat should accrue the hydraulic axial resistance and enforce plants to apply the subsoil water more slowly (Passioura, 1972). An attractive feature of this proposal is that if the soil is damp, then there would be no growth consequence as the nodal root structure, which is very extensive in the upper soil, can effectively supply the crop with water. Under drought stress conditions, several roots morphological traits are modified in response to drought and the morphological modification affects total root length under drought. Small diameter roots enable plants to efficiently enhance hydraulic conductance by increasing the surface area in contact with soil water and the soil volume that can store more water, and small diameter roots also help increase the root hydraulic conductivity by decreasing the apoplastic barrier of water entering the xylem (Comas et al., 2012; Hernández et al., 2010). Consequently, a decrease in root diameter has been suggested as a trait for increasing the plant’s ability to hold water and improve productivity under water deficit (Wasson et al., 2012).

Under drought conditions, roots below 55 cm grow rapidly into lower soil layers to extract soil water from moist soil. The change in root growth in response to drought might indicate drought resistance under water deficit conditions. Under drought conditions, root lengths at soil layers of 55, 75 and 95 cm changed the distribution patterns compared to root lengths under well-irrigated conditions. Root biomass in the deeper soil layers consisted of newly growing roots, elongating roots from old primary roots in the soil layers and branching roots for the roots already existed in the soil layers (Azhiri-Sigari et al., 2000). Under drought conditions, the lateral roots were induced by drying soil in upper soil layers to develop new roots that grow into moist soil in lower soil layers (Nagel et al., 2015). The distribution and architecture of the root structures might depend strongly on the moisture of the deeper soil layer. In this experiment, the difference in irrigated water caused drastic water deficit in the drought treatment and changed the root distribution patterns of sugarcane. Under water stress conditions, the root length of sugarcane in the lower soil layers was higher than under well-watered conditions, and root growth reduces the food that is supplied to shoots.

Underneath early season drought, roots below 55 cm of soil layers have more root tips (root apex zones) than the upper soil layers. The soil moisture in the lower soil layers was higher than permanent wilting point and water was obtainable for the plants. Water stress changed the root structure patterns and increased the root length of the roots below the 55 cm soil layer. Water deficit increases the root length and the percentage of roots in the lower soil layer (Songsri et al., 2008; Jongrungklang et al., 2011). In many of the above studies water stress increased elongation of roots in the deep soil layers, and these previous studies used core sampling method and auger method, which did not recover whole root systems to show the complete root distribution patterns. This behavior suggests that the root responses at given periods of water stress were determined essentially by the root length and changed distribution patterns.

The root system might be a more important sink than the top part of the plant under water deficit at the vegetative stage. The effects of root percentage and the root size might be indicative of a drought resistance mechanism under water-limited conditions. The positive relationship between root length and soil water content by the end of the drought period below the 40 cm soil layer shows the advantage of the increase in deep roots, for extracting water from deep soils over extended periods. Nonetheless, the association between root length and physiological responses to plant water status are very complicated. Several root systems are considered to be essential in sustaining plant productivity under a water deficit. The overall size of root systems is related to the acquisition of water and nutrients from the soil and should be associated with drought resistance and yield performance under drought.

### Physiological traits under different water regimes

Shifts in allometry (metrics of root-to-shoot relationships) and shoot stature can compensate for water shortage and, along with shifts in stand densities, can maintain stomatal conductance under xeric conditions (Mencuccini, 2003; Maseda and Fernandez, 2006). Root length in the deeper soil is an important trait for preserving stomatal conductance under water-limited conditions. The rationale for imposing boundary conditions at the stem base is that the water fluxes through the plant are primarily controlled by stomatal conductance. Indeed, the stomatal conductance is dependent on soil moisture content and relative water content. Although the stomata are fully opened and the stomatal aperture can be visualized in plants that have defective stomatal control and wilt even under well-watered conditions (Borel et al., 2001; Dodd et al., 2009). Under the conditions of water deficit, stomatal closure usually occurs in the afternoon (this can happen even under WW conditions under high evaporative demand). The stomata closure occurs earlier of the day as the soil water reserve is depleted, so stomatal closure in the early morning occurs only at very low soil water potential (Tardieu et al., 1992). Therefore, the soil water potential value that stops water uptake can be interpreted as the triggering stomatal closure and transpiration arrest, even in the early morning.

Studies of plants exposed to drought stress conditions focus on traits, such as root architecture and physiological features (i.e., leaf water potential, osmotic adjustment and RWC) at the vegetative stage (Basu et al., 2016). The C4 plants grown under well-watered conditions and elevated CO_2_ reduced stomatal conductance that can lead to enhanced leaf growth and photosynthesis by mitigating the effects of transient water stress (Seneweera et al., 1998). Root systems of plants responded to soil fertility and soil moisture. Root growth was affected primarily by increased (40–51%) RWC (Derner et al., 2001). Drought-tolerant cultivars of sugarcane maintain a high RWC (Boutraa et al., 2010). The longer roots in the lower soil layers in response to drought are important for plant resistance to drought (Songsri et al., 2008; Jongrungklang et al., 2013). Selection of genotypes with high root length in lower soil layers will likely improve stomatal conductance and enhance photosynthetic capacity and plant growth in a drought-prone environment.

## Conclusions

The sugarcane genotypes displayed different root distribution patterns and architecture. The adaptation of sugarcane subjected to DS condition included a reduction in the root length in the upper soil layer and increased root length in the deeper soil. Differences in the adaptation of the sugarcane genotypes were found for root traits under drought stress, with KK3, MPT06-166, K88-92, CP38-22, Kps01-12 and KPK98-40 showing a high root length in the deeper soil, and this trait could be identified as a drought avoidance mechanism. The present study further revealed that enhanced root length in the deep soil layer is an important trait for maintaining stomatal conductance under drought conditions, and a useful trait for parental selection in future breeding programs as a drought avoidance mechanism.

## Acknowledgments

The authors are grateful for the Thailand Research Fund (TRF) (Grant No. RDG5850007 and RDG5950151) and the Northeast Thailand Cane and Sugar Research Center (NECS), Khon Kaen University (KKU) to providing financial support, and also thankful to the manuscript preparation project of KKU and TRF.

## References

Azhiri-Sigari T, Yamauchi A, Kamoshita A, Wade LJ. 2000. Genotypic variation in response of rainfed lowland rice to drought and rewatering. II. Root growth. Plant Production Science 3, 180–188, DOI:10.1626/pps.3.180

Basu S, Ramegowda V, Kumar A, Pereira A. 2016. Plant adaptation to drought stress [version 1; referees: 3approved]. F1000Research, DOI:10.12688/f1000research.7678.1

Borel C, Frey A, Marion-Poll A, Tardieu F, Simonneau T. 2001. Does engineering abscisic acid biosynthesis in *Nicotiana plumbaginifolia* modify stomatal response to drought?. Plant Cell and Environment 24, 477–489, DOI:10.1046/j.1365-3040.2001. 00698.x

Boutraa T, Akhkha A, Al-Shoaibi AA, Alhejeli AM. 2010. Effect of water stress on growth and water use efficiency (WUE) of some wheat cultivars (*Triticum durum)* grown in Saudi Arabia. Journal of Taibah University for Science 3, 39–48.

Comas LH, Mueller, KE, Taylor LL, Midford PE, Callahan HS, Beerling DJ. 2012. Evolutionary patterns and biogeochemical significance of angiosperm root traits. Journal of Plant Sciences 173, 584–595, DOI:10.1086/665823

Derner JD, Polley HW, Johnson HB, Tischler CR. 2001. Root system response of C4 grass seedlings to CO_2_ and soil water. Plant Soil 231, 97–104.

Dinh HT, Watanable K, Takaragawa H, Kawamitsu Y. 2017. Effects of drought stress at early growth stage on response of sugarcane to different nitrogen application. Sugar Tech DOI:10.1007/s12355-017-0566-y

Dodd IC, Theobald JC, Richer SK, Davies WJ. 2009. Partial phenotypic reversion of ABA-deficient *flacca* tomato (*Solanom lycopersicum*) scions by a wild type root-stock: Normalizing shoot ethylene relations promote leaf area but does not diminish whole plant transpiration rate. Journal of Experimental Botany 60, 4029–4039, DOI:10.1093/jxb/erp236

Doorenbos J, Pruitt WO. 1992. Calculation of Crop Water Requirement. In Crop Water Requirement. FAO of the United Nation, Rome, Italy, pp. 1–65.

FAO. 2016. Food and Agriculture Organization of the United Nations. http://fao.org/faostat/en/#rankings/countries_by_commodity">http://fao.org/faostat/en/#rankings/countries_by_commodity. Accessed 2 October 2018.

Gentile A, Dias LI, Mattos RS, Ferreira TH, Menossi M. 2015. MicroRNAs and drought responses in sugarcane. Frontiers in Plant Science 6, 1–13, DOI:10.3389/fpls.2 015.00058

Hernández EI, Vilagrosa A, Pausas JG, Bellot J. 2010. Morphological traits and water use strategies in seedlings of Mediterranean coexisting species. Plant Ecology 207, 233–244, DOI:org/10.1007/s11258-009-9668-2

Jangpromma N, Thammasirirak S, Jaisil P, Songsri P. 2012. Effects of drought and recovery from drought stress on above ground and root growth, and water use efficiency in sugarcane (*Saccharum officinarum* L.). Australian Journal of Crop Science 6, 1298–1304.

Jongrungklang N, Toomsan B, Vorasoot N, Jogloy S, Boote KJ, Hoogenboom G, Patanothai, A. 2011. Rooting traits of peanut genotype with different yield response to pre-flowering drought stress. Field Crops Research 120, 262–270, DOI:10.1 1/jfcr.2010.10.008

Jongrungklang N, Toomsan B, Vorasoot N, Jogloy S, Boote KJ, Hoogenboom G, Patanothai A. 2013. Drought tolerance mechanisms for yield responses to pre-flowering drought stress of peanut genotypes with different drought tolerant levels. Field Crops Research 144, 34–42, DOI:10.1016/j.fcr.2012.12.017

Kato Y, Okami M. 2011. Root morphology, hydraulic conductivity and plant water relations of high-yielding rice grown under aerobic conditions. Annals of Botany 108, 575–583, DOI:10.1093/aob/mcr184.

Kavar T, Maras M, Kidric M, Sustar-Vozlic J, Meglic V. 2007. Identification of genes involved in the response of leaves of *Phaseolus vulgaris* to drought stress. Molecular Breeding 21, 159–172, DOI:10.1007/s11032-007–9116-8

Maxwell K, Johnson GN. 2000. Chlorophyll fluorescence—a practical guide. Journal of Experimental Botany 51, 659–668.

Maseda PH, Fernandez RJ. 2006. Stay wet or else: three ways in which plants can adjust hydraulically to their environment. Journal of Experimental Botany 57, 3963–3977, DOI:10.1093/jxb/erl127

Mencuccini M. 2003. The ecological significance of long-distance water transport: short-term regulation, long-term acclimation and the hydraulic costs of stature across plant life forms. Plant Cell and Environment 26, 163–182, DOI:10.1046/j.1365-30402 003.00991.x

Nagel KA, Bonnett D, Furbank R, Walter A, Schurr U, Watt M. 2015. Simultaneous effects of leaf irradiance and soil moisture on growth and root system architecture of novel wheat genotypes: implications for phenotyping. Journal of Experimental Botany 66, 5441–5452, DOI:10.1093/jxb/erv290

Neumann G, George TS, Plassard C. 2009. Strategies and methods for studying the rhizosphere—the plant science toolbox. Plant Soil 321, 431–456, DOI:10.1007/ s11104-009-9953-9

Passioura JB. 1972. The effect of root geometry on the yield of wheat growing on stored water. Australian Journal of Agricultural Research 23, 745–752.

Pirnajmedin F, Majidi MM, Gheysari M. 2015. Root and physiological characteristics associated with drought tolerance in Iranian tall fescue. Euphytica 202, 141–155, DOI:10.1007/s10681014-1239–5.

Puangbut D, Jogloy S, Vorasoot N, Craig K. 2018. Root distribution pattern and their contribution in photosynthesis and biomass in Jerusalem artichoke under drought conditions. Pakistan Journal of Botany 50, 879–886.

Robertson MJ, Inmam-Bamber NG, Muchow RC, Wood AW. 1999. Physiology and productivity of sugarcane with early and mid-season water deficit. Field Crops Research 64, 211–227, DOI:10.1016/S0378-4290(99)00042-8

Seneweera SP, Ghannoum O, Conroy J. 1998. High vapour pressure deficit and low soil water availability enhance shoot growth responses of a C4 grass (*Panicum coloratum cv. Bambatsi*) to CO_2_ enrichment. Australian Journal of Plant Physiology 25, 287–292, DOI:10.1071/pp97054

Silva MA, Jifon JL, da Silva JAG, Sharma V. 2007. Use of physiological parameters as fast tools to screen for drought tolerance in sugarcane. Plant Physiology 19, 193–201.

Silva MA, da Silva JAG, Eneiso J, Sharma V, Jifon J. 2008. Yield components as indicators of drought tolerance of sugarcane. Journal of Agricultural Science 65, 620–627.

Smith DM, Inman-Bamber NG, Thorburn PJ. 2005. Growth and function of the sugarcane root system. Field Crops Research 92, 169–183.

Songsri P, Jogloy S, Vorasoot N, Akkasaeng C, Patanothai A, Holbrook CC. 2008. Root distribution of drought-resistant peanut genotypes in response to drought. Journal of Agronomy and Crop Science 194, 92–103,DOI:org/10.1111/j.1439-037x.2008.00296. x

Tardieu F, Zhang J, Katerji N, Bethenod O, Palmer S, Davies WJ. 1992. Xylem ABA controls the stomatal conductance of field-grown maize subjected to soil compaction or soil drying. Plant Cell and Environment.15, 193–197, DOI:10.1111/j.1365-3040.1992. tb01473.x

Thangthong N, Jogloy S, Jongrungklang N, Kvien CK, Pensuk V, Kesmala T, Vorasoot N. 2017. Root distribution patterns of peanut genotypes with different drought resistance levels under early-season drought stress. Journal of Agronomy and Crop Science. 00: 1–12, DOI:10.1111/jac.12249

Thangthong N, Jogloy S, Pensuk V, Kesmala T, Vorasoot N. 2016. Distribution patterns of peanut roots under different durations of early season drought stress. Field Crops Research 198, 40–49, DOI:10.1016/j.fcr.2016.08.019

Unica. 2008. Sugarcane industry in Brazil. http://sugarcane.org/resource-library/books/UNICs%20Institutional%20Folder.pdf. Accessed 27 March 2018.

Wasson AP, Richards RA, Chatrath R, Misra SC, Sai Prasad SV, Rebetzke GJ, Kirkegaard, JA, Christopher J, Watt M. 2012. Traits and selection strategies to improve root systems and water uptake in water-limited wheat crops. Journal of Experimental Botany 63, 3485–3498.

